# Disease-associated mutations in *TPM2* alter regulation of actin filament stability and cofilin-dependent dynamics

**DOI:** 10.64898/2026.05.15.725491

**Authors:** Recep Küçükdogru, Katarzyna Robaszkiewicz, Małgorzata Siatkowska, Joanna Moraczewska

**Author notes:** **Correspondence:** Joanna Moraczewska, Department of Biochemistry and Cell Biology, Faculty of Biological Sciences, Kazimierz Wielki University, Poniatowskiego 12, Str., 85-671 Bydgoszcz, Poland.

## Abstract

Missense mutations in the *TPM2* gene encoding skeletal muscle tropomyosin Tpm2.2 cause congenital myopathies associated with hyper- and hypocontractile phenotypes. Mutation-dependent defects in thin filament stability and length maintenance may contribute to sarcomere dysfunction. To address this possibility, four disease-associated substitutions in Tpm2.2 were analyzed: hypercontractile D20H and E181K, and hypocontractile E41K and N202K. Recombinant proteins were examined *in vitro* for their effects on actin filament polymerization, stability, and cofilin-2-dependent filament length regulation in the absence and presence of troponin (+Ca^2+^).

Wild-type Tpm2.2 inhibited spontaneous actin polymerization and reduced polymerization cooperativity in the presence of cofilin-2. Hypercontractile substitutions D20H and E181K further decreased the polymerization rate, whereas hypocontractile variants had little effect. Under ATP-driven actomyosin interactions, E41K and N202K stabilized filaments, resulting in increased filament length, but this effect was abolished by troponin. All variants slightly decreased cofilin-2 affinity for F-actin without affecting cooperativity. Troponin prevented displacement of Tpm2.2 from the filament at increasing cofilin-2 occupancy, indicating concomitant binding of all proteins to the thin filament, consistent with a structural model based on high-resolution F-actin–Tpm–Tn and cofilactin structures.Tpm2.2-N202K inhibited cofilin-2-dependent depolymerization, whereas Tpm2.2-E181K increased susceptibility to depolymerization. Although cofilin-2 induced filament severing in all cases, the Tpm2.2–Tn complex protected filaments from disassembly.

These findings support a model in which the Tpm2.2–Tn complex forms a cooperative regulatory strand that constrains filament dynamics and transmits structural perturbations along the filament. Disease-causing substitutions differentially alter filament length and stability, potentially contributing to the pathogenesis of myopathies.

## 1 INTRODUCTION

Tropomyosin (Tpm) is a coiled-coil dimer that polymerizes head-to-tail along actin filaments, creating continuous chains on both sides of the filament (Holmes & Lehman, 2008; Lorenz et al, 1995; Pavadai et al, 2020). Tpm2.2, the paralogue encoded by *TPM2*, is ubiquitously expressed in skeletal muscle fibers, where it forms homodimers as well as heterodimers either with Tpm1.1 in fast-twitch fibers or with Tpm3.12 in slow-twitch fibers (Cummins & Perry, 1973; Lehrer & Joseph, 1987; Oe et al, 2007). Each Tpm dimer spans seven consecutive actin subunits and associates with one troponin (Tn) – a three-subunit Ca^2+^-binding complex comprising troponin C (TnC) troponin I (TnI) and troponin T (TnT). Together, Tpm and Tn constitute the core of the thin filament regulatory apparatus, whose classical role is to regulate the interaction between actin and myosin heads in response to Ca^2+^ transients accompanying muscle activation. In essence, the regulatory mechanism involves steric blocking of myosin-binding sites on the actin filament by the Tpm-Tn complex at resting Ca^2+^ levels, followed by the release of this blocked state at activating Ca^2+^ levels, in which myosin cross-bridge cycling is allowed (Brunello & Fusi, 2024; Gordon et al, 2000). However, the functions of the Tpm–Tn complex extend beyond the regulation of actin–myosin interactions. Because the complex spans the entire length of the thin filament, it also has the potential to regulate thin filament length and stability.

Thin filament length varies between muscle types and fiber populations and is tuned to the functional demands of the muscle, thereby contributing to differences in contractile properties (Littlefield & Fowler, 2008; Szikora et al, 2022). Precise control of filament length is therefore essential for proper sarcomere organization and efficient force generation. A protein involved in this process is cofilin-2 (Cof2), the muscle-specific isoform of the actin-depolymerizing factor (ADF)/cofilin family. In muscle fibers, Cof2 localizes to the central region of the sarcomere, near the M-band, where it promotes actin filament disassembly at or near the pointed ends (Kremneva et al, 2014). By regulating filament turnover, Cof2 contributes to the maintenance of thin filament length, and its depletion leads to uncontrolled filament elongation and sarcomere disorganization. Tpm isoforms associated with contractile and cytoskeletal actin filaments have been shown to differentially regulate the severing and depolymerizing activities of Cof2 (Robaszkiewicz et al, 2016; Robaszkiewicz et al, 2023), however, the molecular basis of these differences are not fully understood.

Mutations in human *TPM2* that substitute evolutionarily conserved residues in Tpm2.2 have been associated with a spectrum of congenital muscle diseases (Benarroch et al, 2025; Cassandrini et al, 2017; Laitila & Wallgren-Pettersson, 2021; Tajsharghi et al, 2012). *TPM2* variants have been reported both in congenital myopathies and in arthrogryposes (McAdow et al, 2022; Sung et al, 2003; Tajsharghi et al., 2012). While congenital myopathies are generally characterized by impaired force generation and hypocontractility (Cassandrini et al., 2017; Clarke et al, 2009; Mokbel et al, 2013; Papadimas et al, 2020; Tajsharghi et al, 2007), arthrogryposes result from excessive contractile activity during fetal development, leading to reduced joint mobility (Bamshad et al, 2009; Sung et al., 2003). Since Tpm spans the entire actin filament and controls the access of actin-binding proteins, disease-linked substitutions in Tpm2.2 may perturb the balance between filament stabilization and turnover and, consequently, alter thin filament length regulation, thereby contributing to hypercontractile and hypocontractile phenotypes.

Defining the functional consequences of the mutation provides insight not only into the molecular bases of the corresponding pathologies but also into the mechanisms governing Tpm-dependent regulation of contraction. Recently, we compared effects of four disease-associated missense mutations in *TPM2* on the regulation of actin–myosin interactions (Küçükdogru et al, 2024; Küçükdogru et al, 2025). Amino acid substitutions in Tpm2.2 - D20H and E181K, were linked to hypercontractile phenotypes characteristic of arthrogryposes, whereas E41K and N202K were associated with the hypocontractile states observed in nemaline myopathy. The substitutions were chosen both for their opposing physiological outcomes and for their localization within the N- and C-terminal halves of the actin-binding repeats P1 and P5 (Figure 1A). All substituted amino acid residues are positioned on the solvent-exposed surface of the Tpm coiled coil, rendering them accessible for interactions with proteins that associate with the actin filament. The hypercontractile substitutions are located within regions of Tpm–Tn interaction (Figure 1B). D20H is positioned within the Tpm–TnT junction region (Risi et al, 2023), whereas E181K lies adjacent to the core domain of Tn, comprising all three subunits of the complex (Yamada et al, 2020). In contrast, substitutions associated with hypocontractile phenotypes do not directly interact with Tn (Figure 1B). Our studies demonstrated that hyper- and hypocontractile substitutions differentially perturb actomyosin cross-bridge kinetics (Küçükdogru et al., 2024) and Tn-dependent regulation of contractility (Küçükdogru et al., 2025).

**FIGURE 1.**
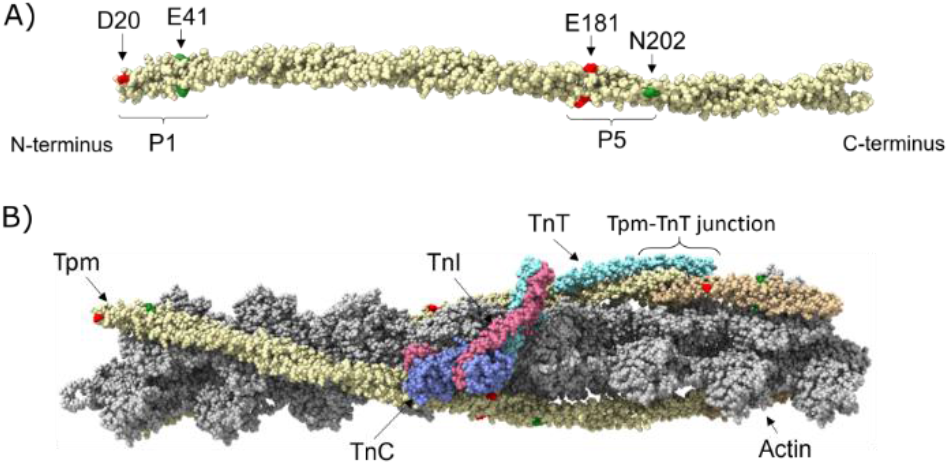
Localization of myopathy-causing substitutions in Tpm2.2. (A) Hypercontractile (red) and hypocontractile (green) substitutions in periods P1 and P5 of tropomyosin (PDB 7UTI). (B) Actin filament in complex with Tpm-Tn with positions of substitutions in Tpm2.2 relative to Tn core domain and Tpm-Tn junction region (PDB: 7UTI). Orientation of the complex shows one Tpm chain bound to Tn core domain and the opposite Tpm chain comprising the Tpm-Tpm overlap region that interacts with N-terminal segment of TnT. For clarity, the second Tn complex is not shown. Actin subunits are shown in silver, Tpm in yellow and gold, TnT in turquoise, TnC in light blue, and TnI in violet. All structural models were visualized using UCSF ChimeraX (Meng et al, 2023).

In this study, we used these Tpm2.2 variants to test the hypothesis that disease-linked mutations affect thin filament stability and Cof2-dependent turnover, both in the absence and presence of Tn. The results indicate that these mutations perturb the balance between filament stabilization and Cof2-mediated turnover, which may contribute to thin filament disorganization in sarcomeres and the associated disease phenotypes.

## 2 RESULTS

### 2.1 Regulation of G-actin polymerization rate and the length of the filament by Tpm2.2 variants

To check whether the disease-causing mutations had an effect on the filament assembly, we compared the polymerization rates of pyrene-labeled G-actin in the absence and presence of Tpm2.2 variants. As expected, the presence of saturating concentration of wild-type Tpm2.2 slowed down actin polymerization (Figure 2A), increasing time required to reach half-maximal polymerization (t_1/2_) by approximately 25%. As indicated by Hill coefficient (*α*^*H*^), Tpm2.2 slightly decreased the cooperativity of the process (Table 1). Compared with wild-type Tpm2.2, the disease-causing amino acid substitutions had no significant effect, except for Tpm2.2-E181K, which further increased t_1/2_, while cooperativity remained unchanged.

**TABLE 1.**
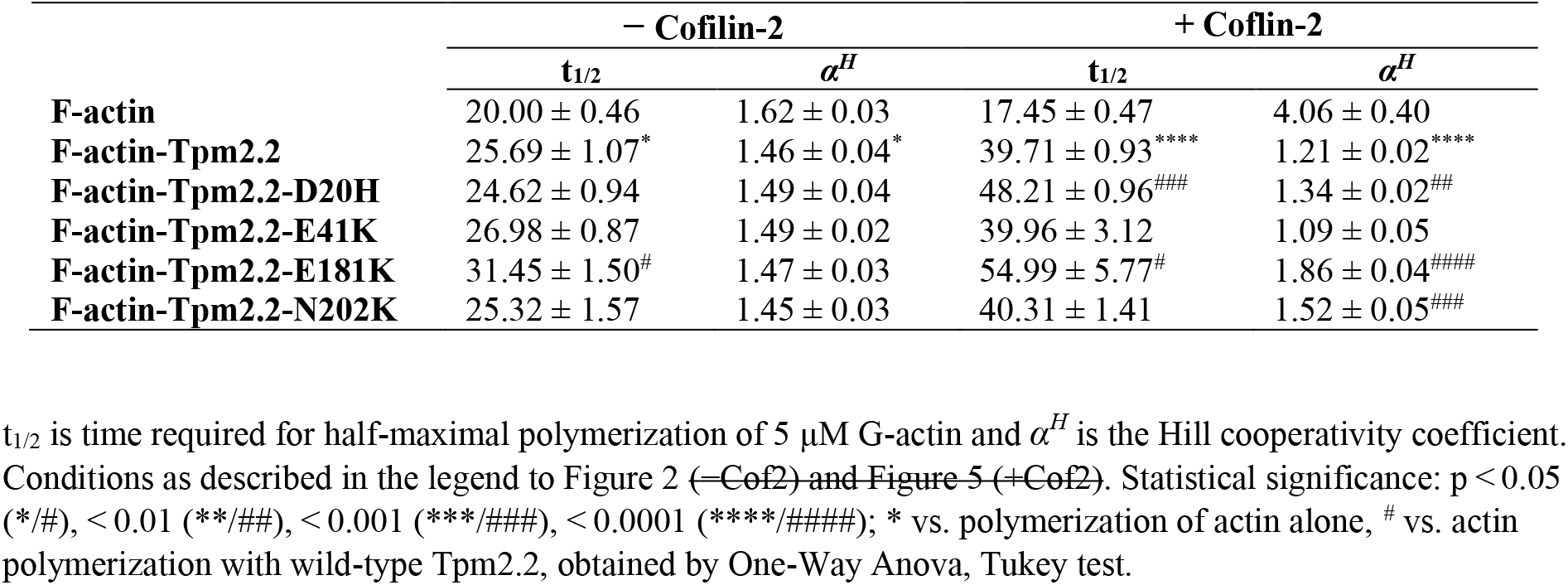
Parameters of G-actin polymerization in the presence of Tpm 2.2 variants.

|  | – Cofilin-2 |  | + Cofilin-2 |  |
| --- | --- | --- | --- | --- |
| | $t_{1/2}$ | $\alpha^H$ | $t_{1/2}$ | $\alpha^H$ |
| **F-actin** | 20.00 ± 0.46 | 1.62 ± 0.03 | 17.45 ± 0.47 | 4.06 ± 0.40 |
| **F-actin-Tpm2.2** | 25.69 ± 1.07* | 1.46 ± 0.04* | 39.71 ± 0.93**** | 1.21 ± 0.02**** |
| **F-actin-Tpm2.2-D20H** | 24.62 ± 0.94 | 1.49 ± 0.04 | 48.21 ± 0.96### | 1.34 ± 0.02## |
| **F-actin-Tpm2.2-E41K** | 26.98 ± 0.87 | 1.49 ± 0.02 | 39.96 ± 3.12 | 1.09 ± 0.05 |
| **F-actin-Tpm2.2-E181K** | 31.45 ± 1.50# | 1.47 ± 0.03 | 54.99 ± 5.77# | 1.86 ± 0.04#### |
| **F-actin-Tpm2.2-N202K** | 25.32 ± 1.57 | 1.45 ± 0.03 | 40.31 ± 1.41 | 1.52 ± 0.05### |

**FIGURE 2.**
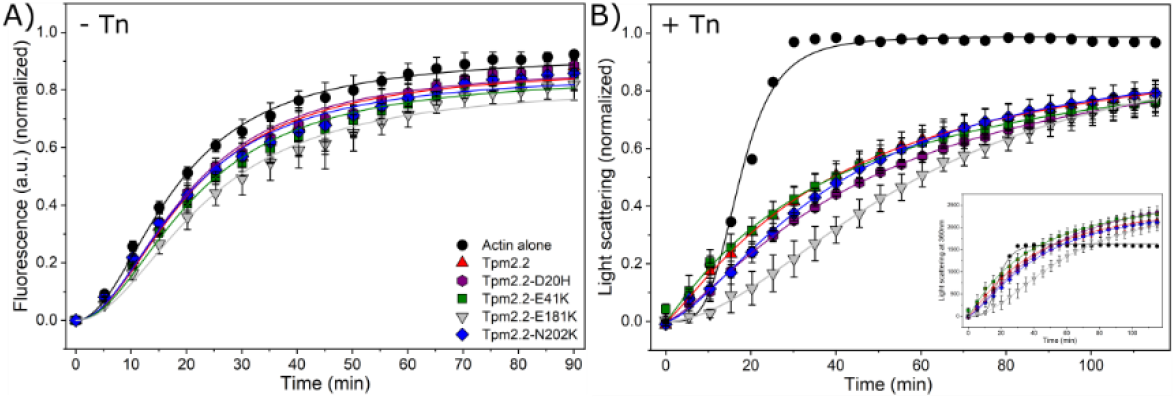
Effects of Tpm2.2 variants on actin polymerization and filament length. (A) Polymerization of pyrene-labeled G-actin monitored by fluorescence in the absence or presence of Tpm2.2 variants. Reactions contained 5 μM actin and 2 μM Tpm2.2 in 20 mM Tris-HCl (pH 7.6), 100 mM KCl, 2 mM MgCl_2_, 0.2 mM ATP, 0.1 mM Ca^2+^, and 1 mM DTT. Data represent the mean ± SEM from 4–5 independent experiments. (B) Polymerization of G-actin in the presence of Cof2 monitored by 90° light scattering in the absence or presence of Tpm2.2 variants. Conditions were as in (A), with 1 μM Cof2 included. Data represent the mean ± SEM from 3–4 independent experiments. (Inset) Polymerization curves normalized to the minimum intensity.

Next, the potential effects of disease-causing mutations on Cof2-dependent actin filament dynamics were examined. As sub-saturating Cof2 concentrations can facilitate severing of formed filaments (Andrianantoandro & Pollard, 2006), a 1:5 Cof2:G-actin molar ratio was used. The rate of polymerization was monitored by 90° light scattering at 360 nm to avoid interference of Cof2 with the fluorescence of pyrene-labeled actin.

The polymerization curves shown in Figure 2B were normalized to the maximal light scattering intensity, demonstrating that, in the absence of Tpm, polymerization was complete within ∼35 min (t_1/2_ = 17.45 ± 0.47), whereas substantially more time was required to reach equilibrium in the presence of Tpm2.2 variants (Table 1). The variants that differed significantly from wild-type Tpm2.2 were the hypercontractile Tpm2.2-D20H and Tpm2.2-E181K, which slowed the initial polymerization rate. Polymerization of unregulated actin was highly cooperative, however, this cooperativity was markedly reduced by all Tpm2.2 variants. Because light scattering intensity is proportional to the mass of polymerized filaments, the higher signal observed in the presence of Tpm2.2 variants (Figure 2B, inset) indicates filament stabilization. No detectable effect of the disease-associated substitutions on the final polymer mass was observed.

In conclusion, Tpm2.2 variants slowed down polymerization rate both in the absence and presence of Cof2, but the final length of the filaments was not affected.

### 2.2 Effects of disease-linked point mutations in *TPM2* on mechanical stability of the actin filaments

To examine filament stability under mechanical load, we analyzed resistance to stress generated by rigor-bound myosin heads and by interactions occurring during the actomyosin cross-bridge cycle. For this purpose, fluorescence microscopy was performed using coverslips coated with HMM, the double-headed heavy meromyosin fragment. Tetramethylrhodamine cadaverine (TRC)-labelled F-actin was used to avoid additional stabilization of filaments by fluorescently labelled phalloidin, which is commonly used for actin visualization in fluorescence microscopy.

First, the actin filaments were attached to myosin heads in the absence of ATP, thereby exposing them to stress generated by rigor-bound myosin. Although filaments appeared longer in the presence of all four Tpm2.2 variants, this difference was not statistically significant (Figure 3A, solid bars). The additional presence of Tn further equalized filament lengths (Figure 3B, solid bars).

**FIGURE 3.**
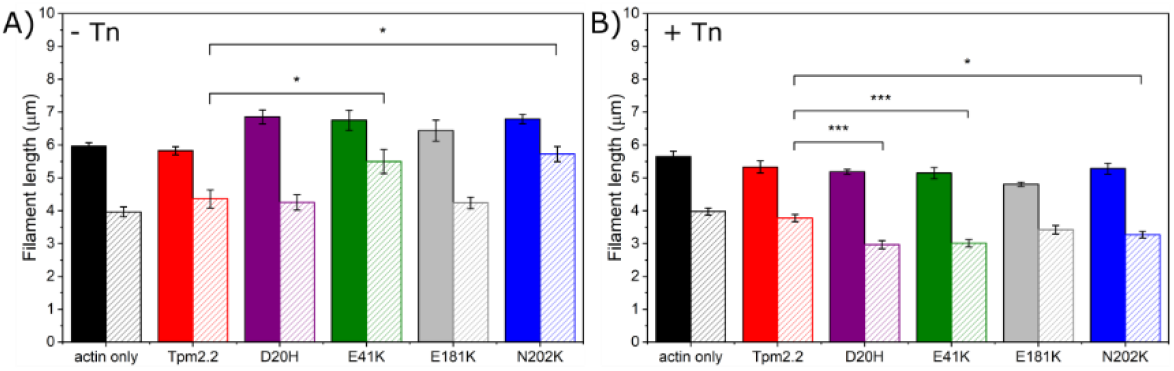
Effects of Tpm2.2 variants on actin filament length under mechanical load. Lengths of F-actin filaments exposed to force generated by myosin HMM in the absence (A) and presence (B) of Tn (+Ca^2+^). Filaments decorated with Tpm2.2 variants were measured before (solid bars) and 30 s after ATP addition (striped bars). Conditions: 0.1 mg/mL heavy meromyosin (HMM), 100 nM F-actin, 34 nM Tpm2.2 variants, and 50 nM Tn in 25 mM MOPS (pH 7.4), 25 mM KCl, 4 mM MgCl_2_, 1 mM EGTA, and 1 mM DTT. Data represent the mean ± SEM from 3–4 independent experiments. Statistical significance was assessed by one-way ANOVA with Bonferroni’s post hoc test (*p < 0.05, ***p < 0.001 vs. wild-type Tpm2.2).

Addition of ATP to the flow cell induced filament sliding along the HMM-coated surface, leading to filament breakage (raw data showing movement of the filaments can be seen in data repository RepOD; the link will be disclosed after publication). After 30 s of movement, all filaments were ∼2 μm shorter than in the rigor state (Figure 3A, striped bars). The hypocontractile variants Tpm2.2-E41K and Tpm2.2-N202K exerted a stabilizing effect, resulting in significantly longer filaments than in the absence of ATP. Filament shortening was also observed when sliding was driven by actomyosin interactions regulated by the Tpm2.2–Tn complex (Figure 3B, striped bars). Under these conditions, the substitutions exerted a destabilizing effect, resulting in significantly shorter filaments. The only exception was Tpm2.2-E181K, for which no statistically significant difference was detected (Figure 3B).

Together, these observations suggest that disease-causing mutations exert a slight but significant destabilizing effect on filaments subjected to the mechanical forces of the actin– myosin cross-bridge cycle.

### 2.3 Effects of disease-linked point mutations in *TPM2* on regulation of Cof2 interactions with the actin filament

Because different Tpm isoforms exert various effects on Cof2 affinity for actin (Robaszkiewicz et al., 2016; Robaszkiewicz et al, 2020), we analyzed the affinity of Cof2 for actin filaments coated with each Tpm2.2 variant using cosedimentation of Cof2 with the thin filament at increasing Cof2 concentrations. The presence of wild-type Tpm2.2 slightly but significantly decreased the affinity of Cof2 for F-actin (Figure 4A). The disease-causing substitutions in Tpm2.2 did not affect significantly Cof2 affinity for actin, except for Tpm2.2-N202K, which showed the weakest ability to reduce Cof2 affinity (Table 2). As assessed from the value of Hill coefficient (α^H^), binding of Cof2 to actin was highly cooperative; however neither of the Tpm2.2 variants affected this function (Table 2).

**TABLE 2.** Parameters of Cof2 binding to actin filaments regulated by Tpm2.2 variants and the troponin complex.

|  | -Tn |  | +Tn (+Ca<sup>2+</sup>) |  |
| --- | --- | --- | --- | --- |
| | K<sub>app</sub> [μM<sup>-1</sup>] | $\alpha^H$ | K<sub>app</sub> [μM<sup>-1</sup>] | $\alpha^H$ |
| **F-actin** | 2.74 ± 0.10 | 4.66 ± 0.83 | | |
| **F-actin-Tpm2.2** | 2.13 ± 0.05** | 5.08 ± 0.61 | 2.33 ± 0.06 | 7.67 ± 1.59 |
| **F-actin-Tpm2.2-D20H** | 2.09 ± 0.04 | 5.47 ± 0.69 | 2.53 ± 0.04<sup>#</sup> | 8.66 ± 1.33 |
| **F-actin-Tpm2.2-E41K** | 2.25 ± 0.14 | 4.05 ± 1.08 | 2.12 ± 0.04<sup>#</sup> | 6.15 ± 0.81 |
| **F-actin-Tpm2.2-E181K** | 2.34 ± 0.11 | 4.25 ± 0.93 | 1.96 ± 0.08<sup>##</sup> | 5.54 ± 1.32 |
| **F-actin-Tpm2.2-N202K** | 2.37 ± 0.07<sup>#</sup> | 5.46 ± 0.92 | 2.28 ± 0.06 | 6.52 ± 0.98 |

K<sub>app</sub> values were obtained from fits to the Hill equation. Statistical significance was assessed using a two-tailed t-test, with $p < 0.05$ considered significant. Symbols indicate significance levels either relative to the F-actin-Tpm2.2 control (<sup>#</sup> $p < 0.05$ , <sup>##</sup> $p < 0.01$ ), or to unregulated F-actin (<sup>\*\*</sup> $p < 0.001$ ).

**FIGURE 4.**
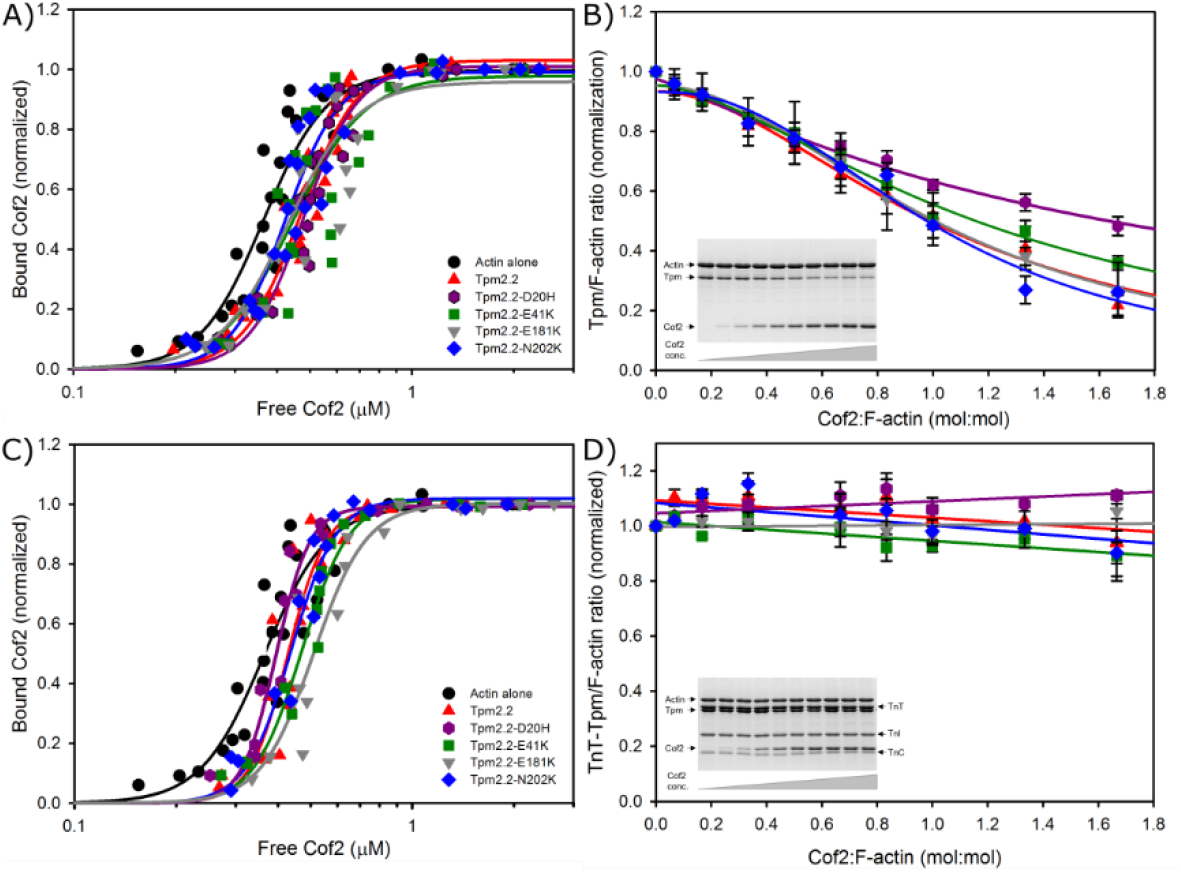
Effects of disease-causing Tpm2.2 substitutions on Cof2 interactions with the thin filament. (**A**) Binding of Cof2 to F-actin in the presence of Tpm2.2 variants. (**B**) Displacement of Tpm2.2 variants from F-actin by increasing of Cof2 concentration. (**C**) Cof2 binding in the presence of Tpm2.2 variants and Tn +Ca^2+^. (**D**) Stability of the Tpm2.2–Tn complex on the filament in the presence of Cof2. Conditions: 3 μM actin, 0.75 μM Tpms and increasing concentration of Cof2 in 5mM Tris-HCl pH 7.6, 100 mM NaCl, 2 mM MgCl_2_, 1 mM DTT. Data represent the mean ± SEM from 3 independent experiments.

Increasing occupancy of the filaments by Cof2 partially displaced Tpm2.2 variants from F-actin (Figure 4B). At saturating Cof2 concentration, only 20–50% of Tpms remained bound to F-actin, with Tpm2.2-D20H being the most resistant to Cof2-induced dissociation (Figure 4B).

In the presence of the Tpm2.2–Tn complex (+Ca^2+^), Cof2 bound to the filament with a similar affinity as in the absence of Tn (Table 2). Due to the scatter of the experimental point, no significant effect of Tn on the cooperativity of Cof2 binding to the thin filament was detected.

In contrast, the effects of the mutations were more variable and resulted in slight but significant changes in Cof2 affinity for actin.

Notably, Tn stabilized all Tpm2.2 variants on the filament and prevented their dissociation by Cof2 (Figure 4D). SDS-PAGE analysis of the pellets showed that Tpm2.2–Tn complexes and Cof2 coexisted on the filaments even at high Cof2 concentrations (Figure 4D, inset), at which Tpm2.2 alone was readily displaced (Figure 4B, inset).

### 2.4 Effects of disease-linked point mutations in *TPM2* on regulation of Cof2-dependent depolymerization of actin filament

We next examined whether the presence of Tpm2.2 or Tn affected the rate of Cof2-mediated thin filament depolymerization. Filament shortening was assessed by directly measuring filament lengths using epifluorescence microscopy. TRC-labeled F-actin, either alone or decorated with Tpm2.2 variants, was immobilized on lipid-covered coverslips and visualized by fluorescence microscopy. This mode of attachment minimized mechanical stress that could influence filament length.

In the absence of Cof2, unregulated filaments ranged from 5.2 to 6.4 μm, with an average length of 5.80 ± 0.52 μm. The average lengths of filaments decorated with wild-type or mutant Tpm2.2 differed only slightly from those of unregulated filaments, and none of these differences was statistically significant. In contrast, myopathy-causing mutations differentially affected filament length in the presence of the Tpm2.2–Tn complex: the hypercontractile variant Tpm2.2-E181K produced shorter filaments (4.3 ± 0.2 μm), whereas the hypocontractile variant Tpm2.2-N202K resulted in longer filaments (7.4 ± 0.4 μm).

After exposure to Cof2, filaments underwent severing, visible as gaps along their length (Figure 5A). Following severing, filaments shortened from both ends, resulting in a substantial reduction in overall filament length. None of the Tpm2.2 variants protected filaments from Cof2-induced severing, as after 30 s fewer Tpm2.2-decorated filaments remained unsevered than unregulated actin (Figure 5B). In contrast, the fraction of unsevered filaments increased to 37% in the presence of wild-type Tpm2.2 and Tn, indicating that the Tpm2.2–Tn complex is a more effective inhibitor of filament severing than Tpm2.2 alone. This protective effect was reduced by all myopathy-causing mutations, although to varying degrees.

**FIGURE 5.**
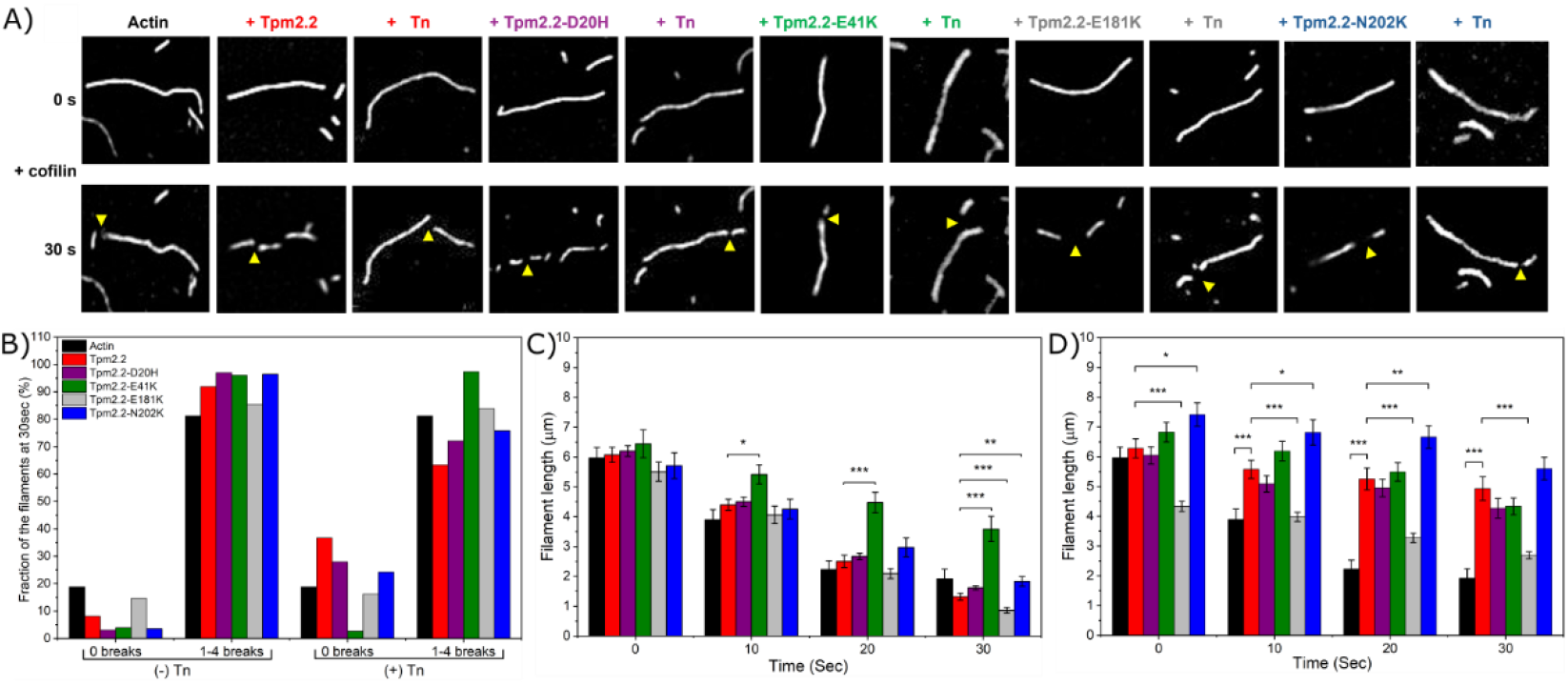
Regulation of Cof2-dependent severing and depolymerization by Tpm2.2 variants. (A) Fluorescence microscopy images of TRC-labeled F-actin (unregulated and Tpm2.2-regulated) before and after 30 s exposure to Cof2. Yellow arrowheads indicate representative severing sites. (B) Fraction of unsevered filaments or filaments containing 1–4 gaps after 30 s exposure to Cof2 in the absence or presence of Tn. (C, D) Cof2-dependent changes in filament length in the absence (C) and presence (D) of Tn. Conditions: 240 nM TRC-labeled thin filaments ± 80 nM Tpm2.2 variants ± 200 nM Tn, 0.1 mM CaCl_2_. Data represent the mean ± SEM from 3 independent experiments. Statistical significance was assessed using Welch’s ANOVA with the Games–Howell post hoc test (*0.01 < p < 0.05, **0.001 < p < 0.01, ***p < 0.001 vs. wild-type Tpm2.2).

Compared with unregulated filaments, the presence of wild-type Tpm2.2 had no effect on the depolymerizing activity of Cof2 (Figure 5C). However, filaments decorated with the hypercontractile Tpm2.2-E181K variant were depolymerized more efficiently. In contrast, the hypocontractile variants, particularly Tpm2.2-E41K, reduced Cof2-dependent depolymerization, resulting in longer filaments than in the presence of wild-type Tpm2.2.

The presence of Tpm2.2-Tn rendered the filaments more resistant to Cof2-mediated depolymerization (Figure 5D). Over the course of depolymerization, the hypercontractile Tpm2.2-E181K produced shorter filaments, whereas filaments coated with the hypocontractile Tpm2.2-N202K remained longer than those with wild-type Tpm2.2.

In conclusion, wild-type Tpm2.2 protects actin filaments from both Cof2-dependent depolymerization and severing only when assembled in complex with Tn, and this stabilization is disrupted in a site-specific manner by myopathy-causing mutations.

## 3 DISCUSSION

Control of thin filament length is a prerequisite crucial for proper sarcomere organization and coordinated muscle contraction. Maintenance of filament length in striated muscle depends on a balance between stabilizing factors, such as the Tpm–Tn complex, and destabilizing activities, including cofilin-mediated turnover. Although key components of this system have been identified (Szikora et al., 2022), the mechanisms by which Tpm-Tn modify the thin filament length and cofilin-dependent actin dynamics, remain incompletely defined. Here, we examined effects of myopathy-linked substitutions located in two functionally important regions of Tpm2.2 (Figure 1) on regulation of the thin filament dynamics. We show that substitutions in Tpm2.2 differentially modulate actin filament stability and Cof2-dependent turnover in the context of the fully regulated thin filament.

### 3.1 Tpm as a regulator of G-actin polymerization

Tpm is known to inhibit spontaneous G-actin polymerization, an effect attributed to stabilization of newly formed filaments through a reduction in the number of elongation-competent filament ends capable of rapid growth (Hitchcock-DeGregori et al, 1988; Wegner & Ruhnau, 1988). In this study, we analyzed unseeded polymerization, which proceeded through nucleation and elongation phases before reaching steady state. Recombinant Tpm2.2 exerted only a modest inhibitory effect on polymerization, which was in contrast to the strong (∼70–80%) inhibition observed for native skeletal muscle tropomyosin and recombinant skeletal Tpm1.1 (Hitchcock-DeGregori et al., 1988; Janco et al, 2016; Robaszkiewicz et al., 2016). These differences may arise from isoform-specific actin affinities of Tpm. However, previous studies have shown that both Tpm1.1 and Tpm2.2 bind actin with similarly high affinity (Janco et al., 2016; Robaszkiewicz et al., 2016; Śliwińska et al, 2021), arguing against an affinity-driven mechanism. Regulation of polymerization rate therefore likely depends on specific interactions between Tpm and actin. This interpretation is supported by our data showing that E181K significantly increased the inhibitory effect of Tpm2.2 on actin polymerization. As demonstrated previously, this mutation does not alter actin affinity but instead changes the orientation of Tpm2.2 on the filament (Küçükdogru et al., 2024). Such subtle differences in binding geometry may change the probability that an incoming actin monomer adopts a productive binding conformation, providing a molecular basis for the regulation of polymerization.

The inhibitory effect of all Tpm2.2 variants on actin polymerization was more pronounced in the presence of Cof2. This effect primarily reflected a reduction in the cooperativity of the process, which in turn decreased the polymerization rate. Cof2 likely destabilizes early actin assemblies slowing down nucleation, while tropomyosin stabilizes and propagates specific filament conformations (Cooper, 2002). Hypercontractile substitutions D20H and E181K were more inhibitory. Since none of the substitutions affected Cof2’s affinity for actin, Tpm2.2 is unlikely to compete with Cof2 for actin or to modify the Cof2–actin interface. Instead, these findings suggest that Tpm2.2 modulates the cooperative behavior of actin filaments in the presence of Cof2, and that the hypercontractile substitutions inhibit the ability of nascent filaments to undergo the conformational adjustments required for productive growth.

### 3.2 Control of thin filament length under mechanical stress

Tpm contributes to the mechanical stability of skeletal muscle thin filaments, which may help maintain filament integrity during repeated actomyosin interactions. This stabilizing effect appears to correlate with the regulation of actomyosin activity, as indicated by lower ATPase activity and slower *in vitro* motility of filaments decorated with hypocontractile Tpm2.2 variants (Küçükdogru et al., 2024). Consistent with this, in the absence of Tn, the hypocontractile substitutions E41K and N202K stabilized actin, resulting in longer filaments.

However, the presence of Tn (+Ca^2+^) introduces an additional layer of regulation. Under these conditions, both hypo- and hypercontractile Tpm2.2 variants produced shorter filaments, despite substantial differences in actomyosin ATPase activity between these groups (Küçükdogru et al., 2025). Actomyosin interactions in the presence of Tn and Ca^2+^ are associated with pronounced changes in filament twisting and bending stiffness, leading to increased flexibility of actin and the generation of internal mechanical stress within the filament (Borovikov et al, 2025). These mechanical effects appear to act as length-equalizing mechanisms that promote filament shortening independently of differences in actomyosin activity among Tpm2.2 variants.

### 3.3 Role of actin-binding period 5 in regulation of thin filament length and Cof2-dependent actin depolymerization

Direct analysis of filament length revealed a notable effect of hyper- and hypocontractile mutations E181K and N202K located in period P5 on Cof2-mediated depolymerization. Both substitutions are located in period 5 (P5), the actin-binding region where Tpm also interacts with the core domain of Tn (Oda et al, 2020; Rynkiewicz et al, 2022; Yamada et al., 2020). While filaments coated with Tpm2.2-E181K-Tn complex were significantly shorter, filaments associated with Tpm2.2-N202K-Tn remained longer than those formed in the presence of wild type Tpm2.2. These effects were not limited to the initial assembly phase but were also maintained under conditions of filament turnover. Since neither Cof2 (this work) nor Tn binding to actin (Küçükdogru et al., 2025) was affected by the mutations, this suggests that the substitutions alter the steady-state of the filament in an allosteric manner. It appears that Tpm2.2, together with Tn, forms a cooperative regulatory strand that stabilizes specific filament states and constrains how structural perturbations, such as those induced by Cof2, are propagated and resolved along the filament. Hyper- and hypocontractile mutations located in P5 do not alter protein binding, but allosterically tune this cooperative behavior, thereby shifting the balance between filament growth, fragmentation, and recovery.

Interestingly, the effects of mutations in P5 on filament length correlated with their associated phenotypes. In skeletal muscle, thin filaments are longer in slow-twitch fibers than in fast-twitch fibers (Gokhin et al., 2012). In this context, the longer filaments observed for the Tpm2.2-N202K variant and the shorter filaments associated with the Tpm2.2-E181K variant are consistent with contractile performance characteristic of hypocontractile and hypercontractile phenotype, respectively. Although filament length in muscle is regulated by additional factors, these observations suggest that mutation-dependent changes in Tpm2.2 function may contribute to phenotype-related differences in thin filament organization. Defective thin filament length regulation may not only impair contractile function but also promote filament aggregation and the formation of nemaline bodies, a hallmark of nemaline myopathy.

### 3.4 Concomitant binding of the Tpm-Tn complex and Cof2 to the thin filament

As demonstrated in this and previous work, Cof2 alters the twist of actin filaments in a manner incompatible with Tpm binding, leading to progressive dissociation of Tpm as Cof2 occupancy on the filament increases (Robaszkiewicz et al., 2016; Robaszkiewicz et al., 2023). However, Cof2 binds to filaments decorated with the Tpm2.2-Tn complex with similar affinity as in the absence of Tn, without displacing Tpm2.2 from the filament. This behavior appears to be independent of the Tpm isoform, as a similar effect was observed for the slow-twitch muscle-specific Tpm3.12 (Robaszkiewicz et al., 2023). Thus, our cosedimentation results indicate that these complexes may co-exist on the F-actin scaffold.

Since high-resolution structure of the F-actin-Tpm-Tn complexes decorated with Cof2 is not available, we used the Matchmaker tool in ChimeraX (Meng et al., 2023), to construct a structural model illustrating the possible concomitant binding of all regulatory proteins on the thin filament (Figure 6). A targeted, chain-specific superposition of corresponding F-actin subunits from the Cof2-bound filament (PDB: 9Y9P) and the Ca^2+^-bound thin filament (PDB: 7UTI) was performed to generate a model predicting simultaneous binding of Tpm–Tn and Cof2 within the same actin filament segment. To show the true helical environment, subunits from the middle of the filament (actin chain A from model 9Y9P and actin chain L from model 7UTI) were aligned. Matchmaker alignment revealed a highly conserved structural core with a pruned backbone RMSD of 1.076 Å, ensuring a high-fidelity alignment. Actin regions comprising four N-terminal amino acids and ten-amino acid sequence in the D-loop (actin subdomain 2) were excluded by the Matchmaker from the structural core because of their high fluctuations in cofilactin. A significant global RMSD of 2.884 Å across all atom pairs highlights a distinct conformational ‘twist’ in the actin monomer induced by Cof2 binding (McGough et al, 1997; Palmer et al, 2026). Steric analysis of this superposed state identified 5,536 atomic clashes at the interface between the Cof2 complex and the regulatory proteins (Tpm, TnT, TnI and TnC). Crucially, these clashes are distributed as broad surface proximities rather than localized steric ‘hotspots’, suggesting that the Ca^2+^-bound ‘Open’ state provides a permissive geometry that can accommodate the Cof2 complex.

**FIGURE 6.**
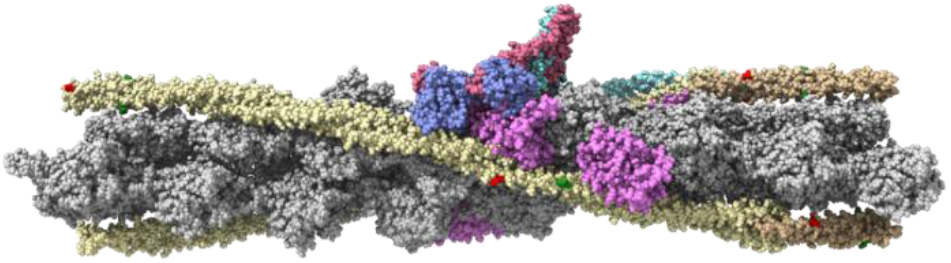
A model of concomitant binding of Cof2 and Tpm-Tn on the actin filament. The model generated in UCSF ChimeraX by aligning the F-actin subunits of the Ca^2+^-bound thin filament (PDB: 7UTI) and the Cof2-bound actin filament (PDB: 9Y9P). Actin is shown in silver, Tpm in yellow and gold, TnT in turquoise, TnC in light blue, TnI in violet, and Cof2 in purple. Red and green show side chains of the substituted amino acids in Tpm. Protein isoforms used to obtain the structures were: α-skeletal actin, cardiac Tpm1.1, cardiac TnT, cardiac TnI, slow skeletal TnC, Cof2 (Palmer et al., 2026; Pavadai et al., 2020). The thin filament model containing Tpm– Tn and Cof2, shown in multiple orientations, and the ChimeraX report are presented in Supplementary Figure S1.

This structural model provides a clear rationale for our biochemical data, suggesting that the activated thin filament does not sterically exclude Cof2, but instead allows for the formation of a hybrid regulatory state where both complexes can co-occupy the actin filament.

## 4 CONCLUSIONS

Missense mutations in the *TPM2* gene, encoding skeletal muscle Tpm2.2, cause congenital myopathies associated with both hypercontractile and hypocontractile phenotypes. Mutations affect thin filament stability and length regulation, which may contribute to severity of the disease. The data support a model in which Tpm2.2, in complex with troponin, forms a cooperative regulatory strand that constrains filament dynamics without altering protein binding. In the presence of troponin, Cof2 no longer displaces Tpm2.2 from the filament, indicating that the regulatory complex remains intact and that the mutation-dependent effects arise from changes in filament behavior rather than protein exchange. Cof2 introduces local structural perturbations, the consequences of which depend on how effectively structural states are stabilized and transmitted along the F-actin-Tpm-Tn complex. These effects are affected by disease-causing mutations and are determined by their localization in Tpm2.2 sequence in relation to the position of Tn on the thin filament.

## 5 MATERIALS AND METHODS

### 5.1 Preparation of native muscle proteins

Actin was purified from the rabbit muscle acetone powder following the method of Spudich and Watt (Spudich & Watt, 1971). After purification, G-actin was stored on ice in G-buffer (5 mM Tris-HCl, pH 7.6, 0.1 mM CaCl_2_, 0.2 mM ATP, 1 mM DTT, 0.02% NaN_3_) and used within two weeks. Protein concentration was determined spectrophotometrically from absorbance at 290 nm using an extinction coefficient of 0.63 for a 0.1% (w/v) protein solution and a molecular weight of 42 kDa.

Heavy meromyosin (HMM) was obtained by limited proteolysis of freshly isolated rabbit skeletal muscle myosin with TLCK-treated α-chymotrypsin according to the procedure described in (Margossian & Lowey, 1982). Molar concentrations of HMM was calculated from absorbance at 280 nm using an extinction coefficient of 0.6 for a 0.1% protein solution, assuming a molecular weight for HMM of 350 kDa. Troponin was isolated from rabbit skeletal muscle using the method of Potter (Potter, 1982). The concentration of Tn was determined from the absorbance at 280 nm using an extinction coefficient of 0.45 for a 0.1% (w/v) protein solution and a molecular weight of 76 kDa.

The fresh rabbit skeletal muscle was a gift from Department of Pathobiochemistry and Clinical Chemistry, Collegium Medicum in Bydgoszcz. The animal handling unit at the Collegium Medicum holds the required approval for procedures involving the sacrifice of animals. Authorization for these procedures was granted to Dr. Piotr Kaczorowski (Consent No. 01/2026) by the Vice-Rector for Collegium Medicum in January 2026, and remains valid until March, 2031).

### 5.2 Expression and purification of wild-type and mutant Tpm2.2

Wild-type and mutant Tpm2.2 recombinant proteins were expressed in *Escherichia coli* (*E*.*coli*) BL21 cells (Thermo Scientific, Carlsbad, CA, USA) and purified as described previously (Küçükdogru et al., 2024). The sequence of the recombinant proteins corresponded to that of human Tpm2.2 and included an Ala–Ser N-terminal extension to compensate for the absence of N-terminal acetylation in bacterially expressed proteins, as described in (Monteiro et al, 1994). The concentration was determined spectrophotometrically using the theoretical molar extinction coefficient of 17,880 M−^1^cm−^1^, calculated with the bioinformatics tool ExPASy ProtParam (https://web.expasy.org/protparam/). Protein concentrations were additionally confirmed by analysis of differential absorption spectra of tyrosine according to (Moraczewska et al, 1999).

### 5.3 Preparation of recombinant Cof2

Recombinant Cof2 was expressed in *E. coli* BL21(DE3) cells transformed with a pGAT2-MM expression vector carrying the mouse Cof2 gene (pPL93; kindly provided by Pekka Lappalainen, University of Helsinki). The construct contained an N-terminal GST tag followed by a thrombin cleavage site.

Protein purification was carried out using GST-affinity chromatography as described previously (Robaszkiewicz et al., 2016). Protein concentration was determined spectrophotometrically at 280 nm using an absorption coefficient 0.79 for a 0.1% protein solution and a molecular weight of 18.5 kDa. Purified protein samples were aliquoted and stored at −80 °C until further use.

### 5.4 Preparation of pyrene-labeled G-actin

Actin was labeled at Cys374 with N-(1-pyrene)iodoacetamide (PIA) (Sigma-Aldrich, Germany) essentially as described previously (Criddle et al, 1985; Kouyama & Mihashi, 1981). G-actin (2 mg·mL−^1^) was polymerized by 0.1 M KCl for 3 h at room temperature, after which PIA was added to a final 1.4-fold molar excess over actin. The reaction mixture was incubated overnight in the dark with gentle stirring. Labeled F-actin was collected by ultracentrifugation, resuspended in G-buffer, and extensively dialyzed to remove excess reagent. Pyrene–G-actin was then ultracentrifuged and separated from oligomers by gel filtration on a Sephacryl® S-200 HR (Cytiva, Germany) column. Protein concentrations and the labeling ratio were determined using the following molar extinction coefficients: 26,600 M^-1^.cm^-1^ at 290 nm for G-actin and 22,000 M^-1^.cm^-1^ at 344 nm for pyrene (Kouyama & Mihashi, 1981).

### 5.5 Analysis of actin polymerization kinetics using fluorescence of pyrene-labeled actin

Actin polymerization was monitored by the increase in fluorescence of 5 μM G-actin containing 5% of pyrene-labeled actin. Actin in G-buffer was polymerized by 100 mM KCl and 2 mM MgCl_2_ either in the absence or presence of 2 μM Tpm2.2 variants. The assay was conducted in a 96-well black microplates (Nunc™ F96 MicroWell™ black polystyrene plate, Thermo Fisher Scientific, USA) at 20 °C. Pyrene fluorescence excited at 366 nm was followed at 387 nm emission wavelength for 90 minutes in a Jasco FP-8350 Spectrofluorometer (Tokyo, JAPAN) equipped with a plate reader. Fluorescence traces were baseline-subtracted and fitted to the Hill equation:

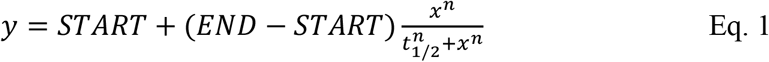

where: y = fluorescence intensity, x = time in minutes; START = minimum fluorescence intensity without polymerization buffer; END = maximum fluorescence intensity with polymerization buffer; t_1/2_ = time required for half maximal polymerization; n = Hill coefficient (α^H^).

To compare the relative potentiation/inhibition across variants, all data were normalized by dividing each experimental point by *END* (maximal intensity) to represent the fractional polymerization. A second fitting to the Hill equation was then performed to determine the t_1/2_ and Hill coefficient (*α*^*H*^).

### 5.6 Analysis of actin polymerization kinetics using light scattering

Light scattering (LS) assay was used to evaluate the effects of Tpm2.2 variants and Cof2 on actin polymerization as described previously (Robaszkiewicz et al, 2021).

For this, 5 μM gel-filtered G-actin, and 2 μM Tpm2.2 variants were mixed with 1 μM Cof2 in G-buffer. Polymerization was induced by addition of 10 × concentrated polymerization buffer to obtain final concentrations of 20 mM Tris-HCl, pH 7.6, 100 mM KCl, 2 mM MgCl_2_, 0.2 mM ATP, 0.1 mM CaCl_2_, and 1 mM DTT. Light scattering was followed immediately for 120 minutes, at 20 °C, and stirring at 500 rpm, in the Jasco FP-8350 Spectrofluorometer (Tokyo, JAPAN) with excitation and emission wavelengths set to 360 nm.

Background signals were subtracted from the raw data, and curves were fitted to a Hill equation (Eq. 1) to determine maximal LS. Datasets were normalized to maximal LS to obtain fractional activity, and Hill equation was used to calculate the apparent half-time (t_1/2_) required to reach 50% polymerization.

### 5.7 Analysis of Cof2 binding to actin filament by co-sedimentation assay

F-actin was prepared by polymerization of 30 μM G-actin in G-buffer (5 mM Tris-HCl, pH 7.6, 0.1 mM CaCl_2_, 0.2 mM ATP and 0.02% NaN_3_) supplemented with 100 mM NaCl and 2 mM MgCl_2_ for 20 minutes at room temperature. The resulting filaments were diluted to 3 μM and incubated with either wild-type or mutant Tpm2.2 (0.75 μM final concentration) in buffer containing 5 mM Tris-HCl, pH 7.5, 100 mM NaCl, 2 mM MgCl_2_, and 1 mM DTT. Stock Cof2 was thawed from −80°C and ultracentrifuged immediately prior to use to remove aggregates. Cof2 was added at concentrations ranging from 0 to 5 μM. Samples were incubated and centrifuged at 40,000 rpm, at 20 °C, for 1 hour using a Beckman Coulter Optima^™^ L-90K ultracentrifuge, rotor type 42.2 Ti. Pellet and supernatant fractions were analyzed by 12% SDS–PAGE.

For experiments performed in the presence of troponin, F-actin (3 μM) was incubated with (0.75 μM Tpm2.2 and 1.8 μM troponin in the presence of 0.1 mM Ca^2+^. Cof2 was added at concentrations ranging from 0 to 5 μM, and samples were processed as described above. Proteins in pellet and supernatant fractions were analyzed by 10% SDS–PAGE.

Band intensities were quantified densitometrically using the ChemiDoc MP Imaging System and Image Lab software v6.1.0.7 (Bio-Rad, Hercules, CA, USA). Ratios of Cof2:F-actin (bound Cof2) and Tpm: F-actin (or Tpm–TnT: F-actin) were calculated from densitometric values and plotted either as a function of unbound Cof2 remaining in the supernatant (Cof2 binding to actin) or Cof2:F-actin molar ratio (dissociation of Tpm/TnT from the filament).

### 5.8 Measurement of actin filament length under mechanical force generated by myosin HMM

The effects of Tpm2.2 variants on filament length during the actomyosin cross-bridge cycle were assessed by fluorescent microscopy using an Olympus IX83 microscope (100 × magnification). G-actin was fluorescently labeled at Gln41 with TRC (Zedira, Darmstadt, Germany) using bacterial transglutaminase, following the protocol established by (Ngo et al, 2016). Briefly, 0.04 mg/mL transglutaminase, 36 μM TRC and 24 μM G-actin (1.5:1 TRC to G-actin molar ratio) were mixed and incubated overnight on ice. Than TRC-G actin was polymerized by overnight incubation with 25 mM KCl, 4 mM MgCl_2_ and used within one week.

Microscopy coverslips were coated with the two-headed proteolytic myosin subfragment (HMM) by immobilizing HMM (0.1 mg/mL) on nitrocellulose-coated coverslips, followed by blocking with 1% (w/v) BSA. A flow-cell was then assembled by attaching the HMM-coated coverslip to a glass slide using parafilm. TRC-F-actin (480 nM) was incubated overnight on ice either alone or with Tpm2.2 variants (160 nM), in the absence or presence of Tn (240 nM) in assay buffer (25 mM MOPS pH 7.4, 25 mM KCl, 4 mM MgCl_2_, 1 mM EGTA, and 1 mM DTT). Non-regulated TRC-labeled F-actin (100 nM) or TRC-labeled F-actin (100 nM) decorated with each of the Tpm2.2 variants (34 nM) and Tn (50 nM) was introduced into the flow cell in assay buffer. Actomyosin interactions were initiated by adding the assay buffer containing 2 mM ATP and 0.4% (w/v) methylcellulose, and oxygen scavenger system (3 mg/mL glucose, 0.1 mg/mL glucose oxidase, and 0.01 mg/mL catalase). Images were recorded for 30 s before and after addition of ATP. Filament lengths were measured using FIJI (ImageJ) software. The average filament length was calculated from all filaments captured across 3 to 4 independent experimental repeats.

### 5.9 Assessment of Tpm2.2 and Cof2-mediated regulation of actin filament length and depolymerization rate

Changes in the filament length were assessed by fluorescence microscopy using F-actin labeled with TRC as described above. TRC-F-actin (480 nM) was incubated overnight on ice either alone or with Tpm2.2 variants (160 nM), in the absence or presence of Tn (240 nM) in assay buffer (25 mM MOPS pH 7.4, 25 mM KCl, 4 mM MgCl_2_, 1 mM EGTA, and 1 mM DTT) with 0.05 % (w/v) BSA. TRC-labeled filaments were introduced in flow cells coated with positively charged lipids (Uchihashi et al, 2012; Yamamoto et al, 2010) and blocked with 10 mg/mL BSA. Unbound filaments were removed with an assay buffer containing an oxygen-scavenging system as described above. The filaments were visualized using an Olympus IX83 microscope (100 × magnification).

To analyze effects of Tpm2.2 variants on Cof2 activity, 240 nM TRC-actin filaments were introduced in the flow cell, followed by 80 nM Tpm variants (±120 nM Tn) and depolymerization was initiated by addition of 240 nM Cof2.

The frames taken in time intervals were analyzed using FIJI (ImageJ) software. The average filament length was calculated from all filaments captured across 3 to 4 independent experimental repeats.

## Abbreviations

Tpm: tropomyosin
Tpm2.2: skeletal muscle tropomyosin isoform 2.2
*TPM2*: gene encoding Tpm2.2
Tn: troponin complex
TnI: troponin I
TnT: troponin T
TnC: troponin C
HMM: heavy meromyosin
S1: myosin subfragment 1
G-actin: monomeric actin
F-actin: filamentous actin
ADF: actin-depolymerizing factor
DTT: 1,4-dithiothreitol
Hepes: 2-[4-(2-hydroxyethyl)piperazin-1-yl]ethanesulfonic acid
MOPS: (3-(N-morpholino)propanesulfonic acid)
ATP: adenosine triphosphate
MW: molecular weight
TLCK: Tosyl-L-lysyl-chloromethane hydrochloride
SDS: sodium dodecyl sulfate
EGTA: ethylene glycol-bis(β-aminoethyl ether)-N,N,N′,N′-tetraacetic acid
BSA: bovine serum albumin
GST: Glutathione S-transferase
PIA: N-(1-pyrene)iodoacetamide
LS: Light scattering
TRC: tetramethyl-rhodamine cadaverine

## AUTHOR CONTRIBUTIONS

*Recep Küçükdogru*: Investigation, Visualization, Formal analysis, Funding acquisition, Writing – review & editing. *Katarzyna Robaszkiewicz*: Investigation, Validation, Writing – review & editing. *Małgorzata Siatkowska*: Investigation, Writing – review & editing. *Joanna Moraczewska*: Conceptualization, Supervision, Writing – original draft, Writing – review & editing, Validation.

## ACKNOWLEDGMENTS

The authors thank Marta Kaczmarek for assistance in actin and recombinant protein preparations. Molecular graphics and analyses performed with UCSF ChimeraX, developed by the Resource for Biocomputing, Visualization, and Informatics at the University of California, San Francisco, with support from National Institutes of Health R01-GM129325 and the Office of Cyber Infrastructure and Computational Biology, National Institute of Allergy and Infectious Diseases. This work was supported by the National Science Centre (NCN), Poland, through the PRELUDIUM 22 funding scheme under grant number 2023/49/N/NZ5/02175.

## DATA AVAILABILITY STATEMENT

The raw data were deposited in the RepOD repository (accession details to be disclosed upon publication).

## AI DISCLOSURE STATEMENT

This paper is an original work and was not prepared using AI, except for language editing. All changes introduced during English refinement were reviewed and approved by the authors.

