## Supplemental Material for "Disease-associated mutations in *TPM2* alter regulation of actin filament stability and cofilin-dependent dynamics"

**A)**

**mmaker #2/A to #1/L**

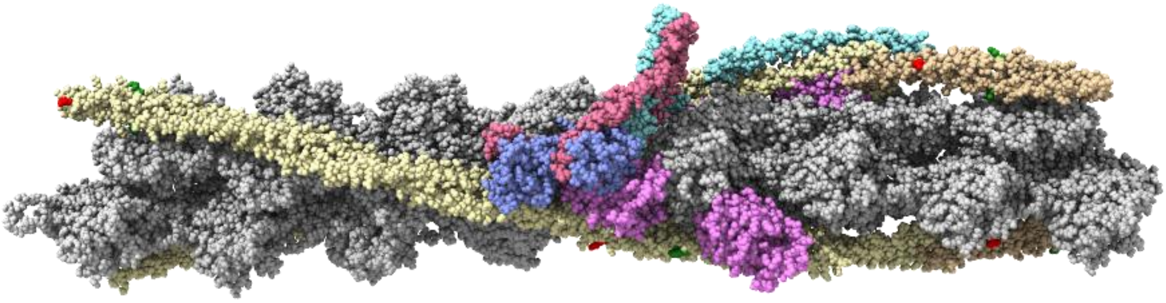

**90 degree turn**

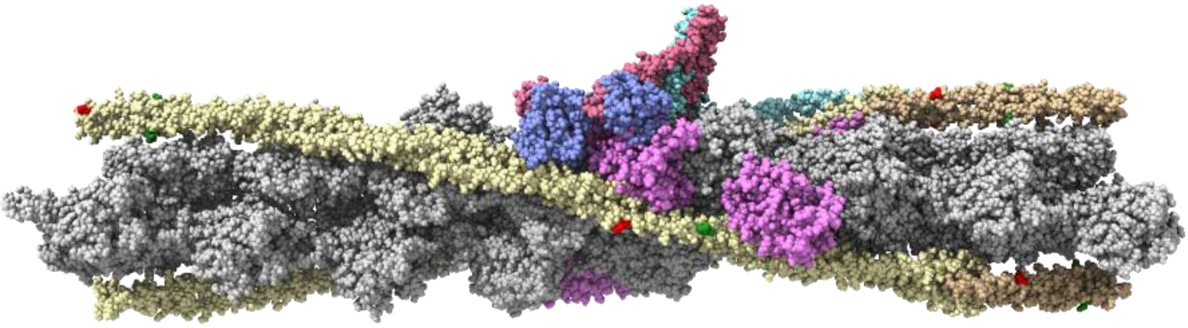

**180 degree turn**

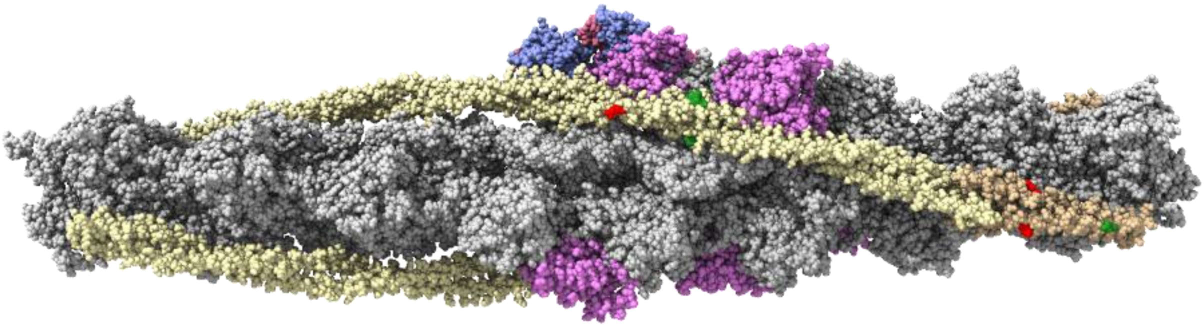

**270 degree turn**

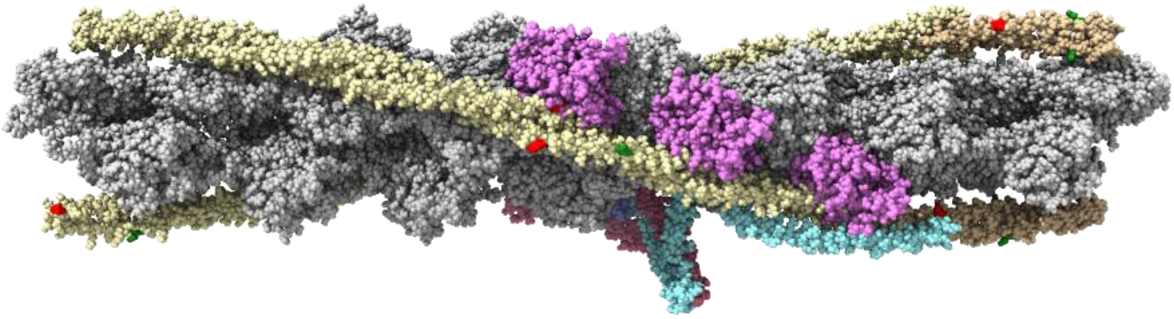

**B)**

### 7UTI, G-actin (chain L) (#1):

- Actin, alpha skeletal muscle (rabbit)
- Cardiac Tropomyosin alpha-1 chain (αα-Tpm1.1) (Human)
- Isoform 6 of Troponin T, cardiac muscle (Human)
- Troponin I, cardiac muscle (Human)
- Troponin C, slow skeletal and cardiac muscles (Human)

### 9Y9P, G-actin (chain A) (#2):

- Actin, alpha skeletal muscle (rabbit)
- Cofilin-2 (Human)

**ChimeraX report:**

Parameters
Chain pairing bb
Alignment algorithm Needleman-Wunsch
Similarity matrix BLOSUM-62
SS fraction 0.3
Gap open (HH/SS/other) 18/18/6
Gap extend 1
SS matrix
H S O
H 6 -9 -6
S 6 -6
At 4
Iteration cutoff 2

Matchmaker 7uti, chain L (#1) with 9Y9P_RK.pdb, chain A (#2), sequence alignment score = 1845.2
RMSD between 221 pruned atom pairs is 1.076 angstroms; (across all 362 pairs: 2.884)

FIGURE S1. A model of F-actin-Tpm-Tn-Cof2. A) The thin filament shown in different orientations. For clarity, only one Tn complex is visualized. Actin is shown in silver, Tpm in yellow and gold, TnT in turquoise, TnC in light blue, TnI in violet, and Cof2 in purple. Red and green show side chains of the substituted amino acids in Tpm. B) Protein isoforms used to obtain the original structures and ChimeraX report on the model building.
